# Protein and DNA Conformational Changes Contribute to Specificity of Cre Recombinase

**DOI:** 10.1101/2024.12.11.627928

**Authors:** Jonathan S. Montgomery, Megan E. Judson, Mark P. Foster

## Abstract

Cre, a conservative site-specific tyrosine recombinase, is a powerful gene editing tool in the laboratory. Expanded applications in human health are hindered by lack of understanding of the mechanism by which Cre selectively binds and recombines its cognate *loxP* sequences. This knowledge is essential for retargeting the enzyme to new sites and for mitigating effects of off-target recombination. Prior studies have suggested that in addition to a few base-specific contacts to cognate *loxP* DNA, the enzyme’s specificity is enhanced by (1) autoinhibition involving a conformational change in the protein’s C-terminal helix, and (2) indirect readout via binding-coupled conformational changes in the target DNA. We used isothermal titration calorimetry (ITC), circular dichroism (CD) and heteronuclear NMR spectroscopy to investigate DNA site recognition by wild-type Cre and a deletion mutant lacking the C-terminal helix. ITC of Cre and a C-terminal deletion variant against cognate and non-cognate DNA recombinase binding elements (RBEs) reveal that the C-terminus reduces DNA binding affinity by six-fold towards cognate DNA. Additionally, ITC revealed highly unfavorable binding enthalpy, which when combined with evidence from CD and NMR of structural differences between cognate and non-cognate complexes support a model in which binding-coupled DNA bending provides a unique structure-thermodynamic signature of cognate complexes. Together, these findings advance our understanding of site-recognition by Cre recombinase.

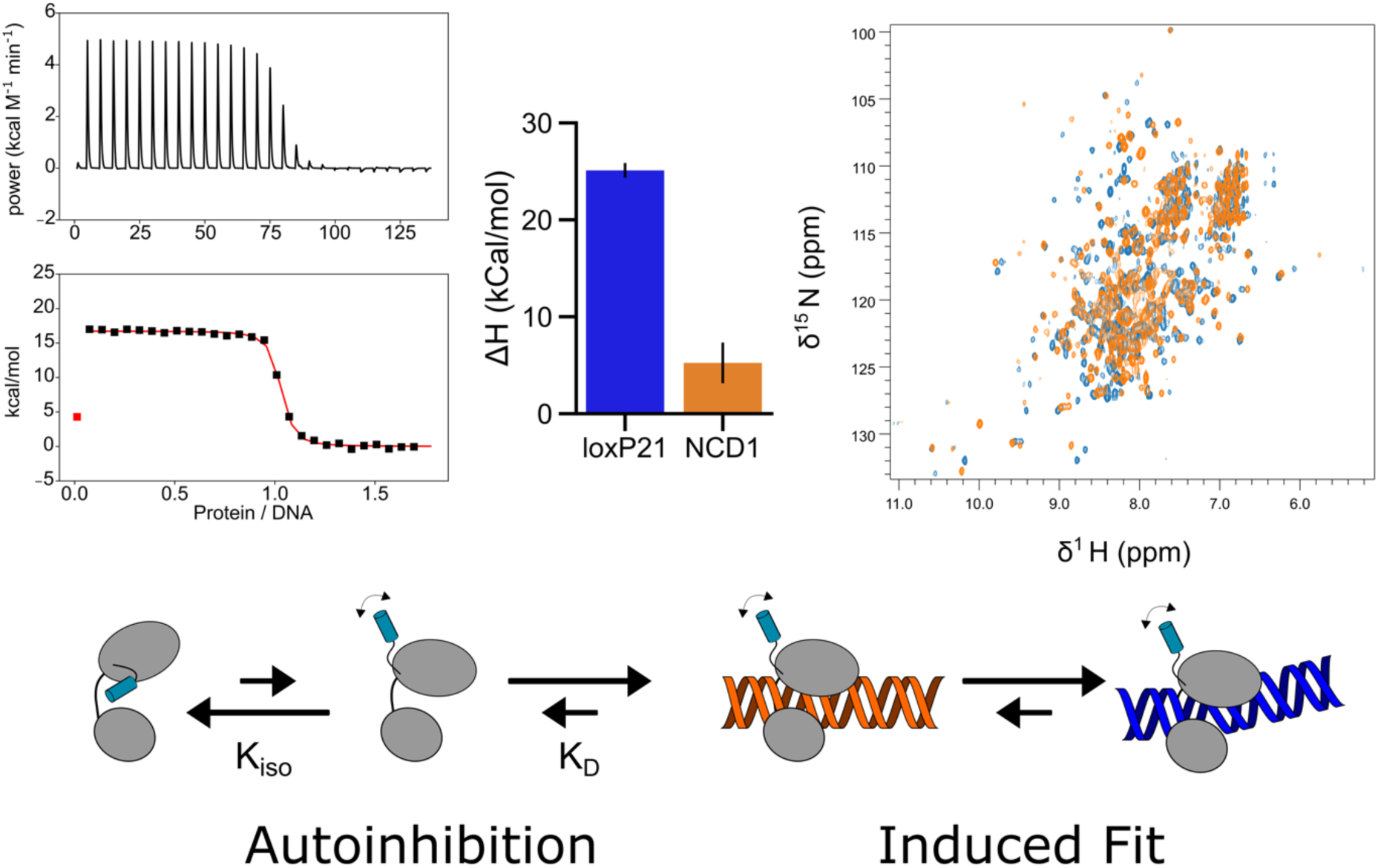

## Introduction

DNA editing technologies present promise toward precision medicine through the targeted manipulation of genetic material^1,2^. Many of these technologies, such as the CRISPR-Cas9 system and zinc finger nucleases, produce site-specific double-strand DNA breaks and rely on host DNA repair systems to incorporate homologous template DNA at these sites^3,4^. However, double-strand breaks can be cytotoxic and are subject to mutagenic host DNA repair by non-homologous end joining and homology-directed repair^1^. Additionally, these tools are not suited for the treatment of diseases resulting from gene transversions^5^. Conservative site-specific tyrosine recombinases (Y-CSSRs) are capable of catalyzing excision, integration, and inversion of targeted DNA sequences without producing double-strand DNA breaks by proceeding through Holliday junction intermediates. They represent a promising complimentary tool in the DNA editing arsenal^6^.

Cre (<u>C</u>auses <u>Re</u>combination) recombinase is a widely used tyrosine site-specific DNA recombinase, employed in laboratory settings to produce transgenic mice and introduce genes through recombinase mediate-cassette exchange technologies^7,8^. Cre recombines *loxP* DNA by forming tetrameric synaptic complexes arranged in an antiparallel fashion (Figure 1). Cre comprises two independently folding domains; an N-terminal core binding domain (Cre^CB^), and a C-terminal catalytic domain (Cre^Cat^)^9,10^. The catalytic domain possesses the essential active site residues, and features a C-terminal sequence (residues 330 to 343) that facilitates trans-docking and cooperative assembly of higher order oligomers, forming helix αN^11–13^.

**Figure 1.**
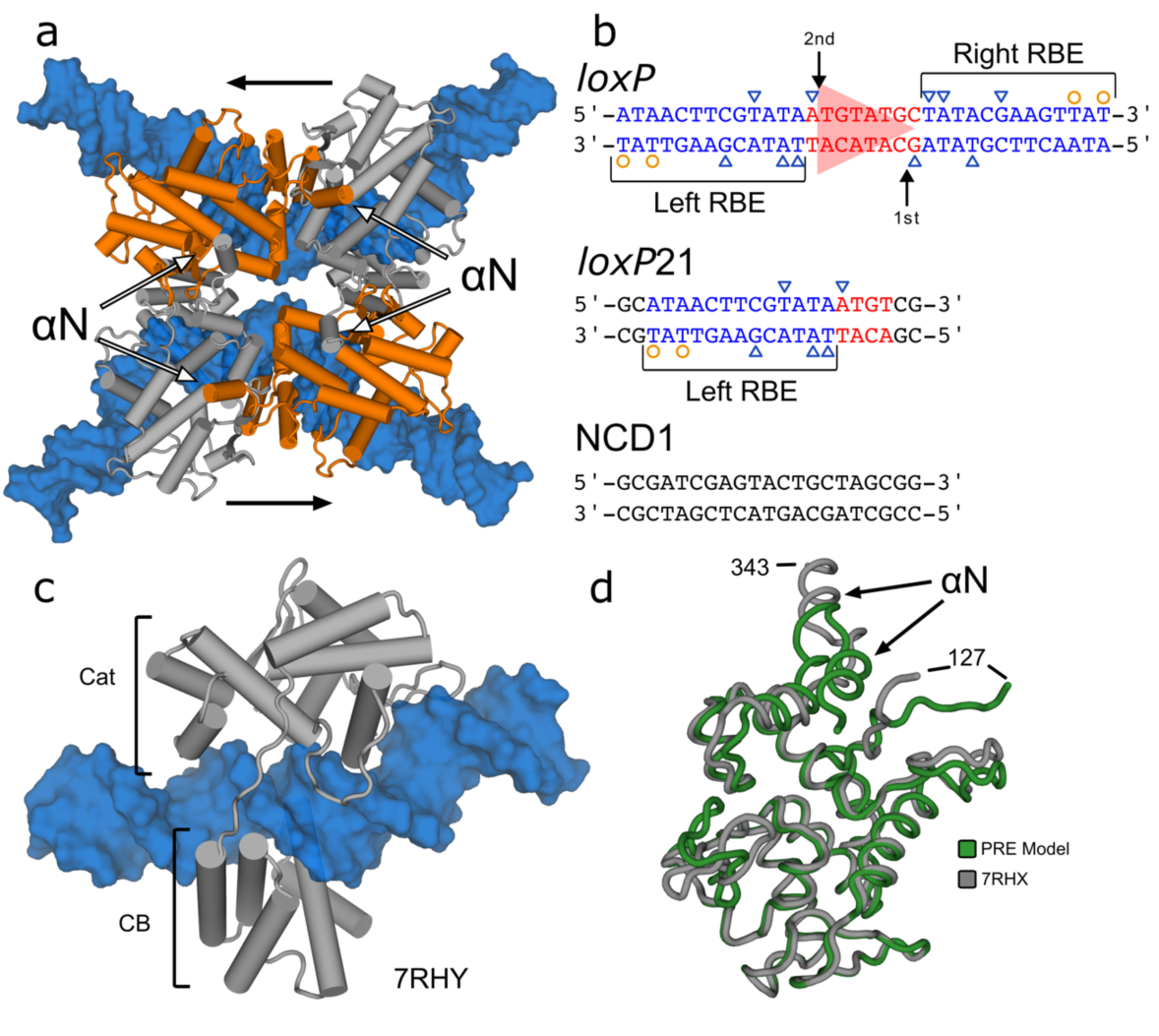
Cre Recombines DNA at *loxP* sites. (a) Cre forms tetrameric Cre_4-_*loxP*_2_ complexes with half-the-sites activity (2HOI) in which alternating pairs of protomers are active for DNA cleavage (orange). Each monomer donates the C-terminal helix into the neighboring protomer either along the DNA substrate or across the synapse. Black arrows indicate the asymmetry in the loxP spacer DNA. (b) Cre binds specifically to *loxP* sequences comprising two pseudo-palindromic recombinase binding elements (RBE, blue) that flank an asymmetric spacer (red). Minor groove contacts are shown as orange circles and major groove contacts are indicated as blue triangles. *loxP*21 is a synthetic *loxP* half-site designed to only allow for monomeric binding and shared homology to the left RBE. NCD1 is a pseudorandom DNA sequence of equal length to loxP21 but share no sequence homology. (c) Cre monomers bind DNA as a C-shape clamp in which both the CB and Cat domains contact the DNA substrate and induce an 18° bend^11^ (7RHY). The C-terminus is not modelled as presumed disorder prevented observable electron density. (d) In the absence of a DNA substrate, Cre adopts an autoinhibited conformation (green) and undergoes a cis-to-trans conformational switch upon DNA binding (grey).

Cre binds with high specificity to *lox*P DNA sequences, comprising two 13 base pair (bp) recombinase binding elements (RBEs), arranged as inverted repeats flanking an eight bp asymmetric spacer (Figure 1). Specificity seems to arise from a combination of direct or water-mediated base contacts, and indirect readout of sequence-dependent structural features of its cognate sites^11,12^. Two Cre protomers bind cooperatively to *loxP* sequences forming Cre_2_-*loxP* dimers, which in turn synapse with another Cre_2_-*loxP* dimer in an antiparallel fashion with respect to the asymmetric spacer to form functional tetrameric (Cre_2_-*loxP*)_2_ complexes^11,14,15^. In the tetrameric complex, Cre exhibits half-the-sites activity, in which only one protomer on each DNA duplex is active for tyrosine-mediated single-strand DNA cleavage, via a mechanism similar to that of type IB topoisomerases^16,17^.

Broader biomedical application of Cre and its derivatives is hindered by a lack of understanding of its specificity determinants. Structural studies of Cre in tetrameric complexes with DNA have identified few base-specific contacts necessary for recombination (Figure 1b), while many non-contacted positions strongly affect recombination activity^18,19^. Additionally, ‘cryptic *lox* sites’ in the human, mouse, and yeast genomes are recombined with efficiencies similar to that of the wild-type *loxP* sequence^20,21^. These observations underscore the need for a more complete understanding of the mechanisms by which Cre achieves specific DNA binding to *loxP* sites and prevents DNA cleavage and recombination of noncognate DNA.

More recent solution NMR and cryo-EM studies suggest two additional sources of DNA-binding specificity: binding-coupled DNA bending, and autoinhibition^9–11^. Crystal structures of Cre in synaptic complexes with *loxP* had revealed large bends in the helical axis, and local groove compression compared to B-form DNA^11,12,15^. Cryo-EM structures of Cre in monomeric and dimeric complexes with DNA revealed progressive DNA bending during the pathway towards synapsis, suggesting that deformability of the DNA sequences could be a significant specificity determinant^11^. In addition NMR studies of Cre in the absence and presence of DNA point to a possible role of autoinhibition: the C-terminal sequence that mediates protein-protein interactions in fully assembled tetramers, docks over the DNA binding interface in the absence of DNA^9^.

In this work, we used isothermal titration calorimetry (ITC), circular dichroism and NMR spectroscopy to characterize Cre-*loxP* site-specific DNA binding. We found that deletion of the C-terminal sequence (331-343) results in a six-fold increase in affinity towards cognate DNA, while a large positive binding enthalpy provides a signature for binding-coupled DNA bending. Circular dichroism (CD) and nuclear magnetic resonance (NMR) spectroscopy corroborated this finding by revealing structural differences between cognate and non-cognate Cre-DNA complexes. Together, these results yield novel insights into the mechanism by which Cre achieves substrate specificity.

## Results

### Autoinhibition by the C-terminus of Cre decreases affinity towards cognate and non-cognate DNA

Previous experiments provided evidence in DNA-free Cre for population of a state in which the C-terminus adopts a conformation predicted to block DNA binding by the catalytic domain^9^. To explore the effect of this conformation on DNA binding, we performed isothermal titration calorimetry (ITC) experiments using full-length Cre and a deletion mutant unable to adopt the autoinhibited conformation, CreΔ330 (1-330) (Figure 2). Solutions containing a *loxP* half site comprising a single RBE (*lox*P21; Figure 1b), or a comparable “non-cognate” DNA with a scrambled sequence (NCD1), were titrated with Cre or CreΔ330. Half-site DNA oligos were selected to prevent cooperative binding of multiple Cre protomers to DNA, thereby monitoring only monomeric DNA binding. Titrating WT Cre into the cell containing *loxP*21 DNA generated large positive heats, indicative of an enthalpically-unfavorable endothermic binding (Figure 2a). Integrated heats were consistent with monomeric 1:1 binding and were fit to a one-site binding model yielding a *K*_D_ of 17.8 ± 2.5 nM, ΔH = +17.1 ± 1.5 kcal mol^-1^, and a corresponding TΔS = 27.7 ± 1.5 kcal mol^-1^ at 15 °C. The same titration using CreΔ330 was also endothermic and following subtractions of heat of dilution, fitting integrated heat results in a *K*_D_ of 2.72 ± 0.84 nM, ΔH of +25.1 ± 0.76 kcal mol^-1^, and TΔS of 36.4 ± 0.8 kcal mol^-1^. These values correspond to a roughly six-fold *increase* in affinity compared to the WT protein despite a 50% less favorable (positive) enthalpy change.

**Figure 2.**
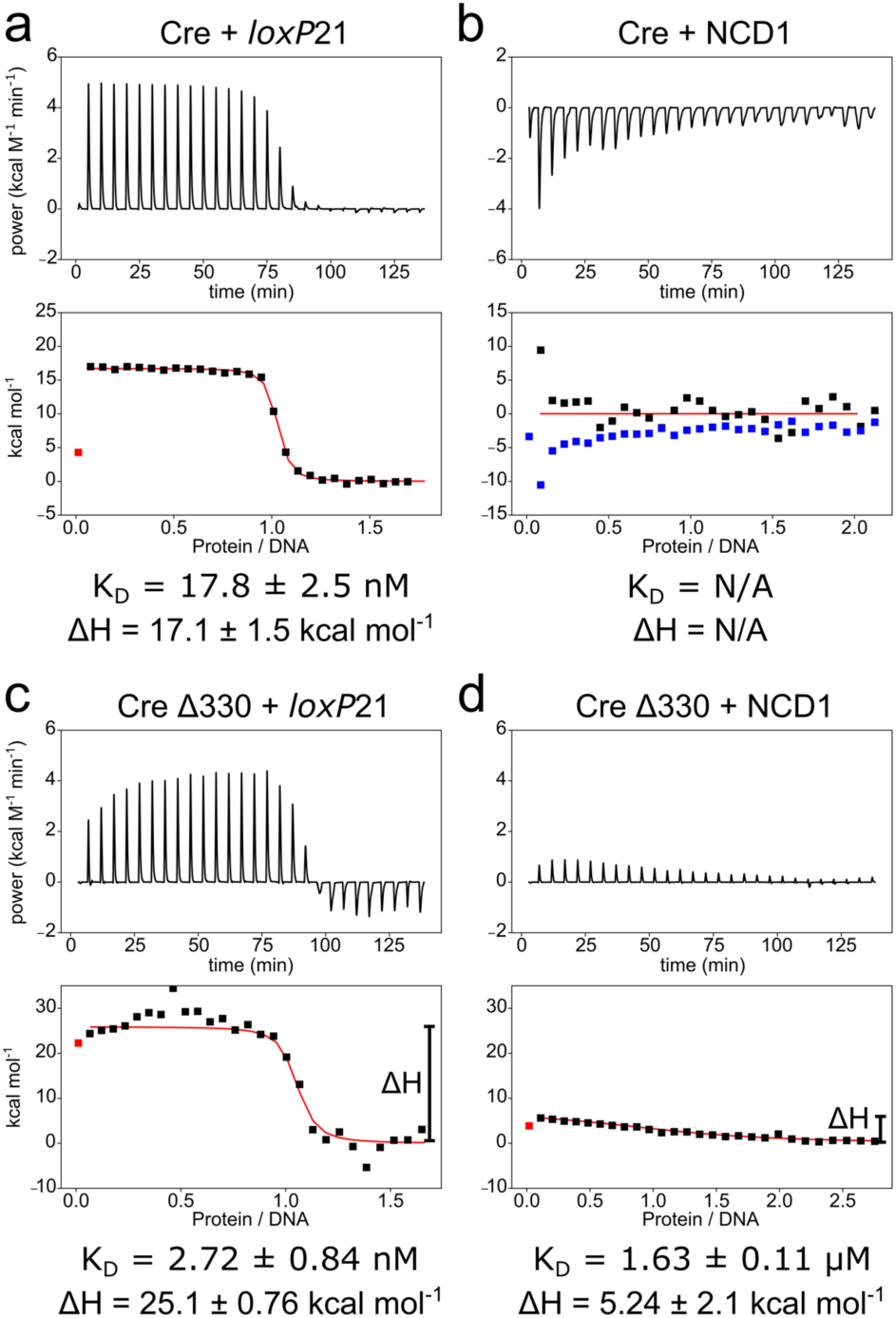
Titration calorimetry reveals autoinhibition by the C-terminus of Cre. Top, thermograms; bottom, integrated heats, for titration of Cre or CreΔ330 into solutions of either cognate (*loxP*21) or non-cognate (NCD1) 21 bp DNA half-sites at 15 °C. In the Cre+NCD1 titration, blue points illustrate heats of dilution that dominate the thermogram. The data indicate that the full-length protein exhibits reduced heat and affinity towards both substrates, while deletion of the autoinhibitory C-terminus results in tighter binding towards both cognate and non-cognate DNA.

We modeled the thermodynamic parameters for Cre binding to a DNA half-site using a three-state model. In this model, a binding-incompetent, autoinhibited state (*I*) exchanges with a binding competent conformation (*P*):

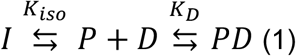

where *K*_D_ is reflects the affinity of the binding-competent conformation of the protein for DNA, and *K*_iso_ = [I]/[P] is the equilibrium constant for isomerization to the autoinhibited conformation. In this model, the effective dissociation constant *K*_D,eff_ for DNA can be described as:

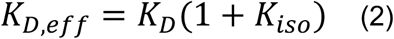

If the experimentally determined DNA binding affinity of CreΔ330 reflects the affinity in the absence of autoinhibition, *K*_D_, and that of full-length Cre corresponds to *K*_D,eff_, we obtain the isomerization equilibrium constant between the two states *K*_iso_ = 5.3 ± 1.7 (equation 2) and a corresponding ΔG_iso_ = -*RT* ln *K*_iso_ = -1.0 ± 0.3 kcal mol^-1^. This corresponds to an autoinhibited population of 85%.

ITC experiments using a pseudo-random non-cognate DNA substrate, NCD1 (Figure 1d) revealed additional features corresponding to autoinhibition. As with the cognate DNA, titrations of NCD1 with CreΔ330 were endothermic, but with lower magnitude (Figure 2d). Fits of the integrated heats to a one-site model yielded a dissociation constant of 1.6 ± 0.1 μM, approximately 600-fold weaker than for binding to cognate DNA, with a ΔH of 5.2 ± 2.1 kcal mol^-1^ and TΔS of -12.9 ± 2.1 kcal mol^-1^ at 15 °C. Given that autoinhibition reduces affinity ∼6-fold towards cognate DNA (ΔΔG = ΔG_ΔαN_ – ΔG_WT_ = -1.1 ± 0.3 kcal mol^-1^), we would also expect WT Cre to exhibit reduced affinity towards non-cognate DNA with a *K*_D_ greater than 10 μM. After correction for heat of dilution, titrations of WT Cre into NCD1 yielded no net heat indicating the interaction is too weak to be measured under these conditions (Figure S1).

The sign and magnitude of ΔH provide interesting clues into the mechanism of DNA site recognition by Cre. Notably, despite binding more tightly to the cognate substrate (*loxP*21), CreΔ330 displays a five-fold less favorable enthalpy of binding compared to that of binding the non-cognate substrate (NCD1). This large unfavorable binding enthalpy is a signature of protein-DNA binding events that induce large deformations in their DNA substrate^22–25^. This observation is consistent with the DNA bending observed in high resolution structures of Cre monomers, dimers and tetramers bound to *loxP* DNA^11,12,15,26^, and with a role for binding-coupled conformational changes in specific binding to binding *lox*P sites. Indeed, sequence-based prediction^27^ of the shape features of the *loxP*21 DNA sequence suggests that it features minor groove compression that is consistent with that observed in the Cre-bound models from cryo-EM and crystallography, while NCD1 is predicted to not exhibit substantial structural variation outside of B-form (Figure S2). These observations motivated additional experiments to examine the role of sequence-specific DNA deformation in specific Cre-*loxP* recognition.

### Cre induces unique deformations in loxP DNA half sites upon binding

To monitor Cre-induced structural changes upon binding DNA, we recorded CD spectra of the free and Cre-bound *loxP21* and NCD1 oligonucleotides in the 250-310 nm wavelength range wherein the protein has no absorbance and reports primarily on base stacking^28^ (Figure 3). For *loxP*21, upon binding Cre we observed large decreases in the molar ellipticity at the 250 and 280 nm minima and maxima, respectively. This change in ellipticity at both 250 and 280 nm is consistent with bending of the DNA substrate^29–31^, as the base stacking interactions that give rise to CD signal are perturbed upon bending of the DNA^28^.

**Figure 3.**
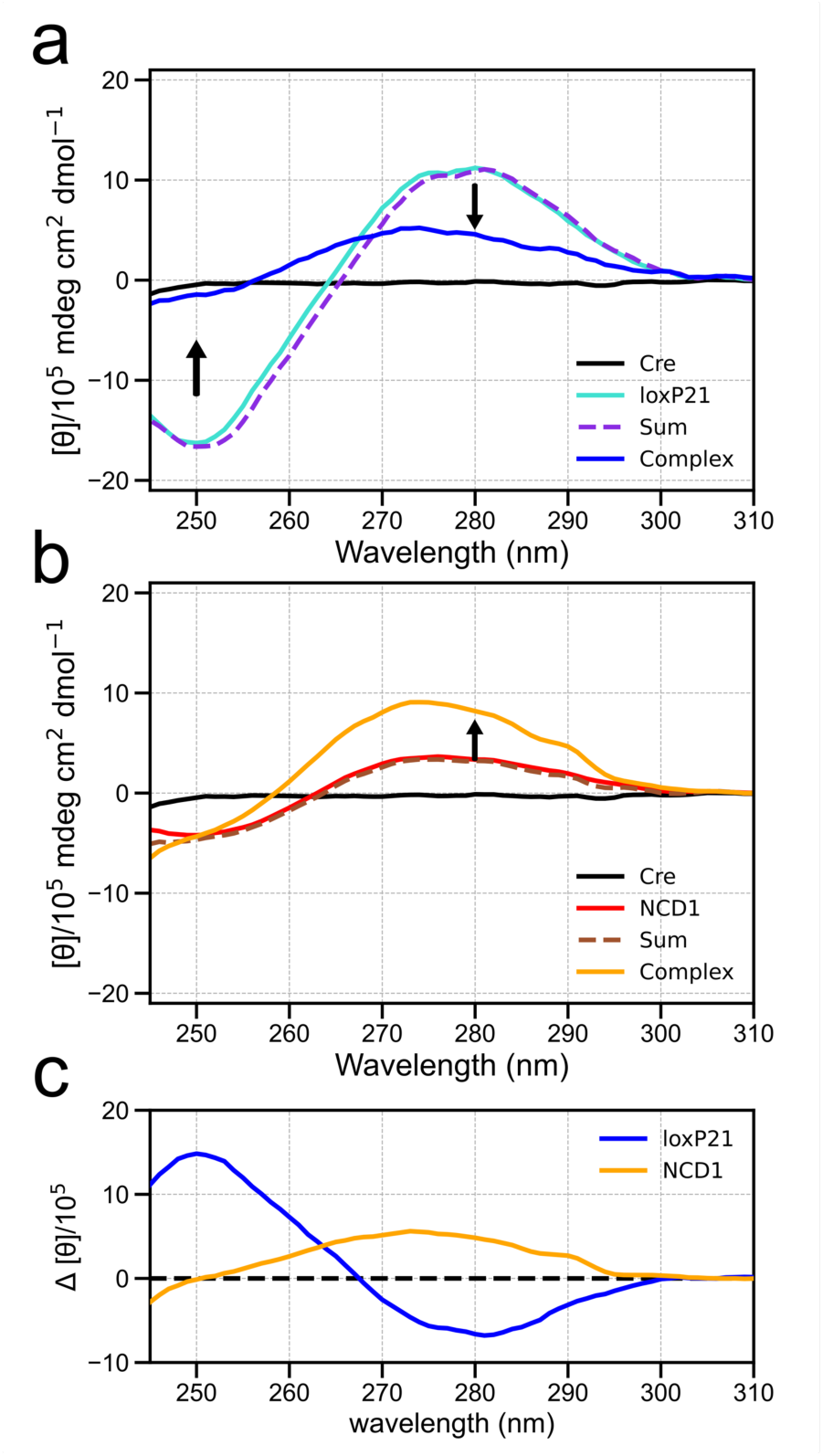
Circular dichroism (CD) reveals distinct Cre-induced distortion upon binding cognate DNA. CD spectra of Cre binding to (a) cognate loxP21 and (b) noncognate NCD1. For loxP21, a large decrease in ellipticity is observed at the 250 nm and 280 nm minima and maxima; for NDC1 ellipticity increases at 280 nm. (c) Differences in CD spectra between the complex and free DNA.

Conversely, binding of Cre to NCD1 (Figure 3b) results in an *increase* in ellipticity at 280 nm and no apparent change at 250 nm, indicating less structural changes and opposite to what is observed for the cognate interaction (Figure 3c). These data suggest that the structural changes in DNA helical structure that accompany Cre binding are distinct between the cognate *loxP* DNA and the non-cognate sequence.

### Cognate DNA Binding Induces Unique Protein Structural Changes

Additional insights into the characteristic signature of site-specific DNA binding by Cre were revealed from two-dimensional ^1^H-^15^N-TROSY-HSQC spectra of [U-^15^N]-Cre bound to either cognate (*loxP*21) or non-cognate (NCD1) DNA. Under the conditions of the NMR experiments, the bound fraction is estimated to be >95% for both *loxP*21 and NCD1. When comparing to the ^1^H-^15^N-TROSY-HSQC spectrum of apo Cre, complexes of both ^15^N Cre-loxP21 and ^15^N Cre-NCD1 show perturbations in chemical shift and similar increased linewidths indicative of binding DNA and tumbling in solution as a larger complex (Figure S3). Additionally, many signals present in the spectrum of free Cre are not observed in the DNA bound complexes, presumably due to large chemical shift perturbation, or excessive line broadening. When overlaying the spectra of Cre bound to loxP21 and to NCD1, widespread chemical shift differences are observed (Figure 4b). Spectra of [U-^15^N] Cre in complex with 1.5 and 2-fold molar equivalents of NCD1 show no significant differences in chemical shifts, indicating the spectrum reflects the NCD1-bound state of Cre (SI Figure 4). These observations indicate that Cre adopts a distinct conformation when bound to to the cognate *loxP* half site in comparison to a noncognate DNA duplex.

**Figure 4.**
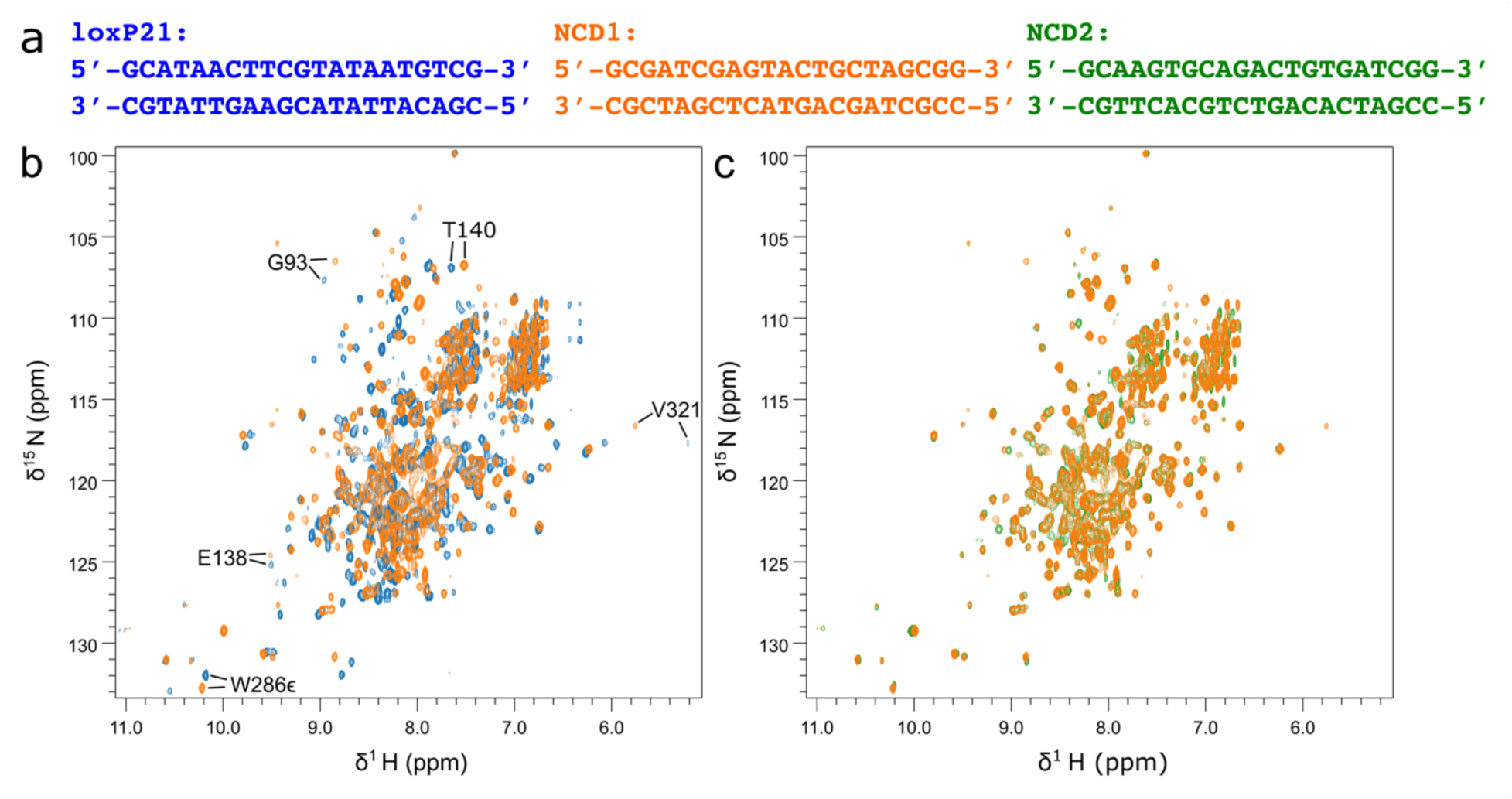
^1^H-^15^N spectra reveal that Cre adopts a unique conformation upon binding *loxP* DNA. (a) Sequences of the loxP21 half site and nonspecific NCD1 and NCD2 dsDNA oligonucleotides. (b) TROSY HSQC spectrum of [U-^15^N]-Cre in complex with either loxP21 (blue) or NCD1 (orange); large spectral differences reflect differences in protein conformation between the two complexes. (c) TROSY HSQC spectrum of [U-^15^N]-Cre in complex with each of the pseudo-random noncognate NCD1 or NCD2 oligos. These spectra overlay quite well, indicating that the protein structure is similar for the two complexes.

Spectra of Cre bound to cognate and non-cognate sequences exhibit distinct features. To test whether the differences in the NMR spectra noted above are dominated by differences in the protein-DNA interface arising from the different DNA sequences, we also recorded spectra of Cre bound to a second non-cognate DNA oligonucleotide, NCD2. Despite limited sequence similarity to NCD1 (Figure 4a), the spectra of the two non-cognate complexes agree exceptionally well, with minimal observed chemical shift differences (Figure 4c). This observation supports the conclusion that binding-coupled conformational changes are major contributors to the large CSPs observed in the cognate Cre-*loxP* complexes.

While the spectra exhibited substantial peak broadening consistent with the overall reduced tumbling rates for the >50 kDa Cre-DNA complexes, well-resolved signals could be tentatively assigned for direct comparison. Notably, Cre-*loxP* complexes exhibit chemical shift perturbations for amides distal to the DNA-binding interface, such as G93, T140, and V304, whereas these signals are largely unperturbed from their positions in the apo-Cre in the noncognate complexes (Figure 5a). These data indicate unique conformational changes occur when Cre binds to *loxP* sequences that do not occur when binding noncognate DNA.

**Figure 5.**
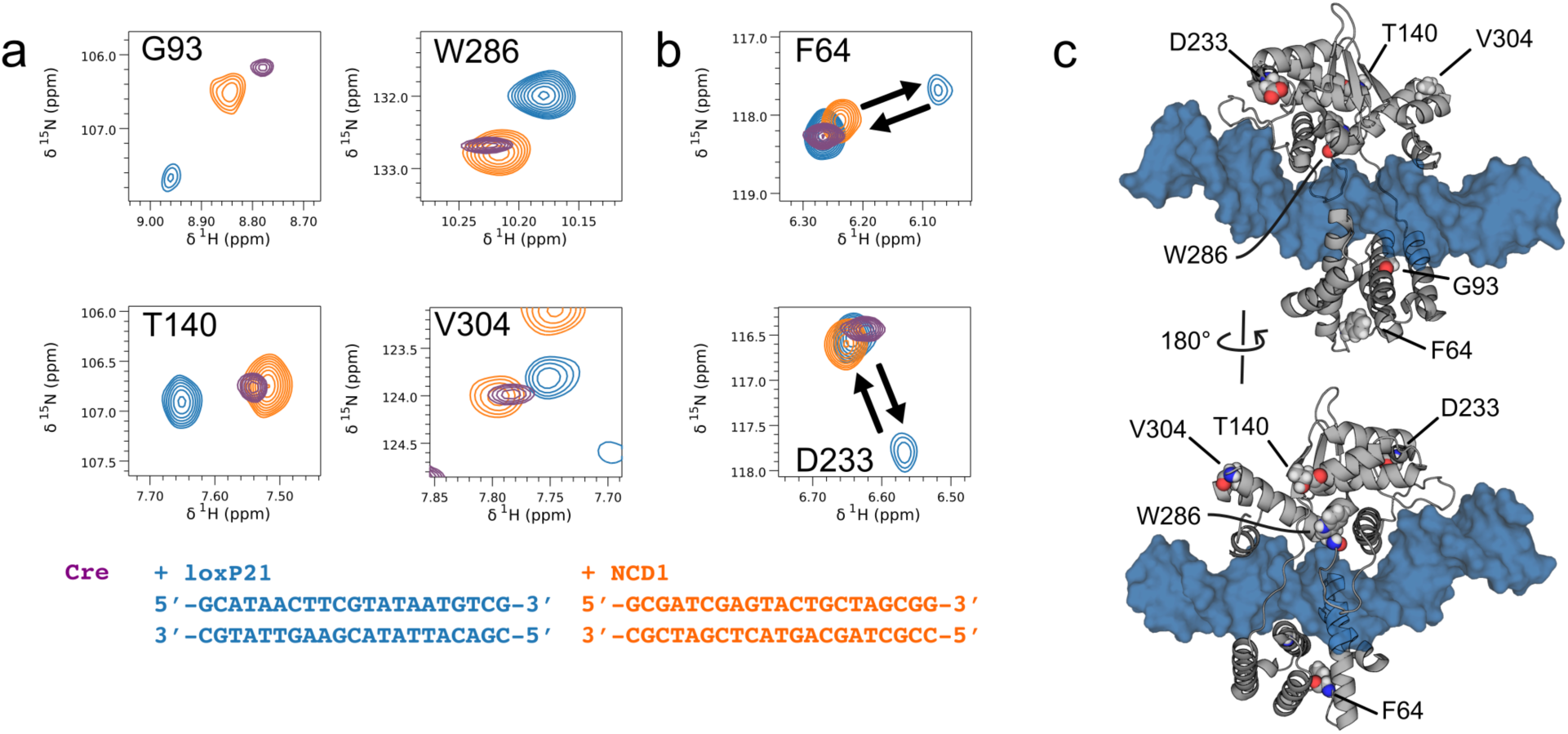
Cognate Cre-*loxP* complexes feature distinct NMR spectral signatures. (a) Regions of the NMR spectra of Cre in the absence (Apo, purple), and presence of cognate (blue) and noncognate (orange) DNA. (b) Doubling of some signals in the αB-C loop (F64) and the αI-J loop (D233) reveal emergence of slow-exchange dynamics in the cognate complex that is not observed in non-cognate complexes. (c) Model of Cre monomer bound to a *loxP* half-site illustrates that many CSPs in the cognate complex occur away from the DNA binding interface, indicative of coupled conformational changes.

In addition to widespread chemical shift differences between cognate and noncognate complexes, spectra of Cre bound to cognate DNA reveal changes in slow time-scale dynamics as evident by peak doubling. This is evident from signals from Phe64 and Asp233 which both lie away from the DNA binding interface (Figure 5b). Cre possesses nine proline residues distributed across the core binding and catalytic domains, and residues adjacent to these prolines appear to exhibit peak-doubling only when bound to cognate, *loxP* sequences. Notably, peak doubling is not observed in either the apo protein or the non-cognate complexes, indicating that this is a feature of site-specific recognition and may have consequences for subsequent assembly of higher order complexes.

## Discussion

We used biophysical approaches to test the hypothesis that autoinhibition and unique conformational changes enhance DNA binding specificity of Cre recombinase. Titration calorimetry illustrated that the C-terminus of Cre inhibits binding to DNA substrates and revealed an unfavorable (positive) binding enthalpy that hinted at an induced fit mechanism for site recognition. Spectroscopic measurements by CD and NMR provided strong support for binding-coupled conformational changes consistent with an indirect readout mechanism in sequence-selective DNA binding by Cre. These observations provide valuable insights into the mechanism of site selection by Cre.

### A role for autoinhibition in site-selection by Cre

Autoinhibition has been observed in other sequence-specific DNA binding proteins, wherein adopting a binding-incompetent conformation decreases affinity towards DNA substrates^32–35^. Examples include the architectural HMGB1 protein, which contains a negatively changed C-terminal tail that interacts with the DNA binding surface^33^, and the winged helix-turn-helix protein ETS-1, whose flanking helices similarly occlude its DNA binding surface^35^. For Cre, structural evidence for the population of an autoinhibited conformation involving the C-terminus was provided by (1) NMR chemical shift perturbations upon its deletion, and (2) paramagnetic relaxation NMR experiments which showed that the labeled C-terminus is proximal to its DNA binding surface^9^. Indeed, here we report that deletion of the C-terminal helix of Cre results in ∼6-fold increase in binding affinity to *loxP* sites, in agreement an autoinhibition model (Equations 1 and 2). Likewise, calorimetric titrations using noncognate DNA show that presence of the C-terminal sequence strongly reduces Cre affinity towards noncognate DNA sequences compared to *loxP* half sites.

What might be the functional consequences of autoinhibition of DNA binding by Cre? The N-terminal CB and C-terminal catalytic domains of Cre fold independently, while autoinhibition by the C-terminal peptide inhibits DNA binding by the C-terminal catalytic domain. This configuration would potentially allow relatively weak DNA binding by the CB domain and transient sampling of a DNA bound configuration by the Cat domain. This weakly bound conformation, with both fast association and dissociation rates, could facilitate rapid scanning of long DNA substrates by exchange between transient weak and long-lived high affinity complexes with DNA^36,37^. Such a mechanism has been previously proposed to facilitate sequence scanning and site selection in other DNA binding proteins, including the tandem Zn-finger domains of Egr-1 and the architectural protein HMGB-1^32,38,39^. Considering that Cre has been shown to recombine *loxP* DNA sequences separated by many kilobases in eukaryotic DNA, it seems likely that the ability to sample low-affinity, fast-scanning conformations, in addition to high-affinity, long-lived complexes, is an important contributor to that activity.

### A role for induced-fit in site-selection by Cre

The presented calorimetric and spectroscopic experiments provide strong evidence for a major role of indirect readout in site recognition by Cre. Binding of Cre to *loxP* half-sites incurs an enthalpic penalty ΔH of +17 kcal mol^-1^, which is countered by a favorable increase in entropy TΔS of 28 kcal mol^-1^ at 15 °C (Figure 2). Binding of full-length Cre to non-cognate DNA generated insufficient heats for accurate analysis, but CreΔ330 bound the same oligo with a five-fold decrease in unfavorable ΔH. Using machine learning models trained on known structures of protein-DNA complexes^27^, the *loxP* sequence is predicted to show localized groove narrowing and distortion from the typical B-DNA structure. This behavior is unique when compared to the NCD1 sequence (Figure S2). Structures of Cre bound to half-site DNA by Cryo-EM^11^ (7RHY) and crystallography^40^ reveal the same pattern of protein-DNA hydrogen bond and ion-pair interactions as observed in tetrameric complexes, in addition to local deviations in DNA structure from canonical B-form. CD spectra of the *loxP* and non-cognate half-site recorded in the absence and presence of Cre (Figure 3), reveal large and opposite changes in molar ellipticity at wavelengths sensitive to changes in base stacking. Based on these observations we conclude that the large unfavorable enthalpy of Cre binding to *loxP* half-sites arises in large part from the energetic cost of deforming the *loxP* DNA.

Cognate *loxP* DNA half-site recognition is also coupled to unique structural changes in Cre. Two-dimensional ^1^H-^15^N correlated NMR spectra of Cre bound to cognate and noncognate DNA reveal site-specific structural perturbations to the protein (Figure 4, Figure S3). DNA binding to either cognate or non-cognate DNA half-sites generates widespread chemical shift perturbations, many of which can be attributed to their direct proximity to DNA. More informative are shift perturbations far from the DNA binding interface, because these reveal differences in protein structure between cognate and non-cognate complexes. Indeed, for many of the resolved signals from residues distal to the DNA interface, chemical shifts from noncognate complexes closely resemble those of the apo protein, whereas they adopt distinct chemical shifts in the cognate complex (Figure 5). These observations suggest that upon cognate sequence recognition, mutual binding-coupled conformational changes occur in both protein and DNA substrate.

The findings here using *loxP* half-site substrates can be expected to extend to site selection in the context of recognition of full *loxP* sites by Cre. Compared to half-site complexes, structural studies of Cre bound as a dimer to the *loxP* inverted repeats reveal additional conformational changes, including more extensive DNA bending and formation of ordered protein-protein interfaces^11^. These interactions lead to cooperative binding to DNA and contribute to *loxP* recognition^12,41–43^, while specific recognition of the first Cre protomer to one half-site is likely essential for effectively recruiting the next protomer in a Cre_2_-*loxP* assembly.

### Implications for application of Cre-loxP technology

Previous studies have used substrate-linked directed evolution to derive mutants with altered specificity and expand Cre’s use from that of genetic engineering in the laboratory towards the improvement of human health^5,44,45^. However, the existing framework for how Cre achieves specificity towards *loxP* sequences was incomplete. Previous work studying Cre-*loxP* specificity focused on recombination efficiency between *loxP* and mutant lox sequences^18,46,47^. Indeed, mutations in *loxP* that significantly reduce recombination efficiency are also predicted to perturb DNA minor groove width at positions coincident with bending in Cre-*loxP* half sites^18,46^ (Figure S6). Presumably, these mutations also reduce cooperative assembly of dimers and the stability of the synaptic interface stabilizing tetrameric complexes. While the work presented herein compares thermodynamics of Cre binding *loxP* half-sites and pseudo-random noncognate sequences, we observe the same trend in which predicted DNA shape elements are indicative of the experimentally observed indirect readout mechanism driving DNA binding specificity. Thus, our conclusions drawn from thermodynamic measurements should be consistent and transferable to *loxP* mutants with observed reduction in recombination efficiency.

In summary, the presence of an autoinhibited conformation dynamically blocks the DNA binding interface of the catalytic domain and may also play a role in the ability for Cre to rapidly scan DNA and locate *loxP* sequences. Additionally, the lack of base-specific contacts between Cre and *loxP* appears to be overcome through an indirect readout mechanism, in which bending of *loxP* sequences and coupled conformational changes in Cre produces a stable cognate complex compared to noncognate DNA. Future work will be necessary to understand the functional role of autoinhibition in DNA scanning and how sequence elements in *loxP* allow for this indirect-readout mechanism. Together, our findings enhance our understanding of how Cre maintains specificity towards *loxP* DNA and provides an improved framework for engineering new specificity and identifying promising DNA targets for Cre-mediated recombination.

## Methods

### Protein Expression and Purification

The pET21A plasmid encoding wild-type (WT) Cre (P06956) was provided by Gregory Van Duyne (University of Pennsylvania Philadelphia, PA). The CreΔ330 truncation mutant was constructed by introducing a TAG stop codon following residue 330 using the Q5 Site directed mutagenesis kit (NEB) and primers 5’-CCTGGATAGTtagACAGGGGCAATG-3’ and 5’-TTACGGATATAGTTCATGAC-3’. Plasmid vectors encoding WT and Cre ΔN were used to transform electrocompetent *E. coli* T7 express LysY/Iq cells (New England Biolabs) by electroporation in a 1 mm cuvette and plated on LB agar plates containing 0.1 mg/mL carbenicillin and chloramphenicol for antibiotic selection. Proteins were overexpressed by growing cells in 1 L of either LB media or M9 minimal media (9.465 g Na_2_HPO_4_, 0.18 g KH_2_PO_4_, 4 g NaCl, 2 mM MgSO_4_, 10 mL MEM Vitamin Solution (Gibco), 4 g glucose, and 1 mL of trace metal mixture: 50 mM FeCl_3_, 20 mM CaCl_2_, 10 mM MnCl_2_, 10 mM ZnSO_4_, 2 mM CoCl_2_, 2 mM CuCl_2_, 2 mM NiCl_2_, 2 mM Na_2_MoO_4_, 2mM Na_2_SeO_3_, 2 mM H_2_BO_3_) [U-^15^N] supplemented with 1 g/L ^15^NH_4_Cl (Cambridge Isotopes) as the sole nitrogen source and 0.1 mg/mL carbenicillin and incubated at 37 °C, shaking at 220 rpm. Once the cultures reached sufficient optical density at 600 nm (0.6-0.8) measured using a 1 cm cuvette, cells were induced with 400 μL of 1M IPTG (Isopropyl β-D-1-thiogalactopyranoside) and incubated for an additional 4 hours at 37 °C shaking at 220 rpm. Following the induction period, cells were pelleted by centrifugation at 4225 x g and pellets were frozen at -80 °C for storage.

Frozen pellets were thawed on ice and resuspended in buffer containing 40 mM Tris pH 7, 100 mM NaCl, 1 mM EDTA, one tablet of miniComplete EDTA free protease inhibitor (Roche) and 17.5 μM PMSF (phenylmethylsulphonyl fluoride). Cells were lysed by sonication on ice (Qsonica, 50% amplitude for 6 minutes, 5 seconds on and 5seconds off), and the insoluble fraction was pelleted by centrifugation at 26891 x g for 45 minutes at 4 °C. The resulting supernatant was filtered using a 0.2 μm filter (Pall) and first purified using cation exchange chromatography using a 5mL SPFF column (Cytiva) with protein eluted using a 75 mL linear gradient (40 mM Tris pH 7.0, 0.1-1 M NaCl). Fractions containing Cre were isolated and diluted (1:4) in low salt buffer (40 mM Tris pH 7.0, 0.1 M NaCl) and further purified by affinity chromatography using a 5 mL heparin column (Cytiva) with a 110 mL linear gradient (40mM Tris pH 7.0, 0.1-1.5 M NaCl). A final purification step was done using size exclusion chromatography with a 120 mL S75pg column (Cytiva) in buffer containing 40mM Tris pH 7.0, 500 mM NaCl, and 5 mM DTT. Cre elutes as a single peak, and final purity was ensured to be >95% by SDS-PAGE.

### DNA Purification

Lyophilized single strand DNA oligos were obtained from Integrated DNA Technologies and resuspended in miliQ H_2_O to a concentration on 1 mM measured by nanodrop (Thermo Scientific). Double-stranded DNA was produced by mixing equal amounts of top and bottom strands, heating in a 95°C water bath for 10 minutes, and annealing by cooling slowly overnight. Single-strand DNA contamination was removed from annealed duplexes by anion exchange chromatography using a QHP column (Cytiva) with a linear elution gradient (10mM TRIS pH 7.0, 100 mM to 1 M NaCl) and purity was ensured by 2% agarose gel supplemented with 0.1% (v/v) Sybr Safe dye (Invitrogen) for visualization.

### Isothermal Titration Calorimetry

Purified WT Cre, CreΔ300, and DNA substrates were dialyzed overnight against ITC buffer containing 20mM HEPES pH 7, 250 mM NaCl in either 6-8 kDa MWCO dialysis tubing (Spectrum Laboratories) or 3.5 kDa MWCO dialysis cassettes (Thermo Scientific). Titrations were performed on a Microcal VP-ITC platform (Malvern Panalytical). Macromolecule concentrations were determined by measuring absorbance at either 260 or 280 nm using a 1cm quartz cuvette (ε_280,WT_ = 48,930 M^-1^ cm^-1^, ε_280,Δ330_ = 49,180 M^-1^ cm^-1^, ε_260,loxP21_ = 333,780 M^-1^ cm^-1^, ε_260,NCD1_ = 338,155 M^-1^ cm^-1^,). Protein concentration in the syringe were 20-150 μM and DNA concentration in the cell were 2-15 μM. ITC experiments were carried out at 15 °C with a reference power of 5 μCal/s. Protein solutions were injected at volumes of 10 μL with an inter-injection delay of 300 seconds and a stir speed of 307 rpm. Baselining and integration of each injection was completed using NITPIC^48^ and titrations were fit using ITCSIMLIB to a single-site binding model.^49^

### Circular Dichroism

Purified WT Cre, *loxP*21, and NCD1 were dialyzed against CD Buffer (20mM HEPES pH 7, 100mM NaCl) overnight at 4 °C. Free Cre, *loxP*21, and NCD1 concentrations were determined by measuring absorbance at 280nm and 260nm for protein and DNA respectively and samples were prepared with concentrations of 5 μM, 5 μM, and 8 μM for Cre, *loxP*21 and NCD1 respectively. For *loxP*21 complexes, equal volumes of 10 μM Cre and *loxP*21 were mixed and transferred to a 1 mm cuvette. For NCD1 complexes, Cre and NCD1 were mixed at volumes necessary to yield a final concentration of 50 μM Cre and 25 μM NCD1 following concentrating to 1 mL. The mixture was then dialyzed into CD buffer overnight at 4 °C and concentrated to 1 mL for analysis. Following data collection, the concentration of each component was determined by measuring absorbance at 260 and 280 nm and calculating concentration using extinction coefficients of each species at 260 and 280 nm (ε_280,Cre_ = 48,930 M^-1^ cm^-1^, ε_260,Cre_ = 25,229 M^-1^ cm^-1^, ε_280,NCD1_ = 182,083 M^-1^ cm^-1^, ε_260,NCD1_ = 338,155 M^-1^ cm^-1^). Final concentrations of the components were determined to be 35 μM Cre and 15 μM NCD1. CD spectra were recorded using a JASCO J-1500 CD with a 1 mm quartz cuvette at room temperature. Spectra were scanned from 350 to 190 nm at a scanning speed of 100 nm/min and digital integration time of 2 s. Each spectrum was signal averaged with 5-10 accumulations.

### Nuclear Magnetic Resonance

NMR samples of free Cre were produced by dialyzing purified Cre into buffer containing 20mM Hepes pH 7.0, 100 mM NaCl and 5 mM DTT overnight at 4 °C. Following dialysis, Cre was concentrated using a 10 kDa cutoff spin filter to 100 μM, with concentration measured by nanodrop (Thermo Scientific). Cre-DNA complexes were assembled by slowly mixing protein into either cognate (*loxP*21) or noncognate (NCD1 or NCD2) DNA at high ionic strength (500 mM NaCl) to a final molar ratio of 1:1.5 protein:DNA. The complexes were then dialyzed stepwise (300 mM ➔ 100 mM ➔ 50 mM ➔ 25 mM NaCl, each 100-fold dilution and for >8 hours at 4 °C) into NMR buffer containing 20 mM HEPES at pH 7, 25 mM NaCl, and 5 mM DTT in a 3.5kDa MWCO dialysis cassette (Thermo Scientific). Dialyzed protein-DNA complexes were then concentrated using a 3.5 kDa cutoff spin filter to approximately 200 μM based on protein and DNA concentrations prior to mixing. Samples of 500 μL of complex supplemented with 10% (v/v) D_2_O and 225 μM DSS (2,2-dimethyl-2-silapentane-5-sulfonate) as an internal standard were added to 5 mm NMR tubes. ^1^H-^15^N TROSY-HSQC spectra of the complexes were recorded using a Bruker Avance 800 MHz spectrometer equipped with a triple resonance inverse (TXI) cryoprobe. Spectra were recorded using 40 scans comprising 2048x280 complex points and a spectral width of 18.026 x 50.01 ppm centered at 4.70 and 117.0 ppm in the ^1^H and ^15^N dimensions. Data was processed and analyzed using NMRFx Analyst^50^ (nmrfx.org).

## Supporting information

Supporting information: additional supplemental figures comprising 15N-1H NMR spectra, simulated ITC data, and predicted DNA shape plots (PDF).

## Supporting Information

Supporting information: additional supplemental figures comprising ^15^N-^1^H NMR spectra, simulated ITC data, and predicted DNA shape plots (PDF).

## Acknowledgements

Dr. Chunhua Yuan of the Ohio State Central Campus Instrument Center (CCIC) for assisted with NMR experiments, and Dr. Alicia Friedman of the OSU Biophysical Characterization of Interactions Facility (BCIF) supported CD and ITC instrumentation. Foster Lab members assisted with sample preparation, data collection, and fruitful discussion, particularly Caroline Hervey and Justin Perdomo. This work was supported by NIH R01 GM122432 (to M.P.F). and J.S.M. was supported by NIH T32 GM144293.

