## Supporting information: additional supplemental figures comprising 15N-1H NMR spectra, simulated ITC data, and predicted DNA shape plots (PDF). for "Protein and DNA Conformational Changes Contribute to Specificity of Cre Recombinase"

$$K_{iso} = \frac{[I]}{[P]} \quad and \quad K_a = \frac{[PD]}{[P][D]}$$

$$K_{eff} = \frac{[PD]}{([I] + [P])([D])} = \frac{[PD]}{([P]K_{iso} + [P])([D])} = \frac{[PD]}{[P][D](1 + K_{iso})}$$

$$K_{eff} = \frac{K_a}{1 + K_{iso}}$$

$$K_{D,eff} = K_D(1 + Kiso)$$

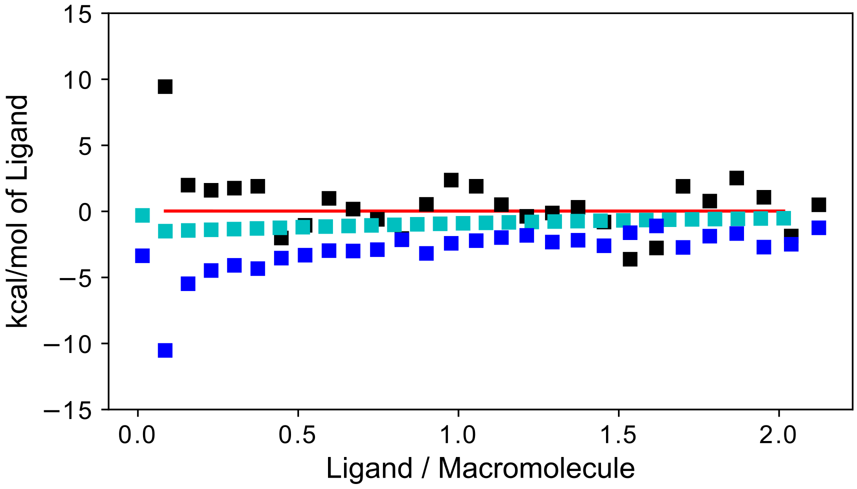


**Supplemental Figure 1.** ITC titration of Cre into NCD1. Heats of dilution (blue) were subtracted from integrated heats for each injection to yield binding enthalpies (black) fit to a one site model (red). Using our model that suggests a $K_{iso}$ of ~5.5 and observed $\Delta H_{iso}$ of 8.0 kCal/mol we simulated the binding curve for Cre binding to NCD1 (cyan).

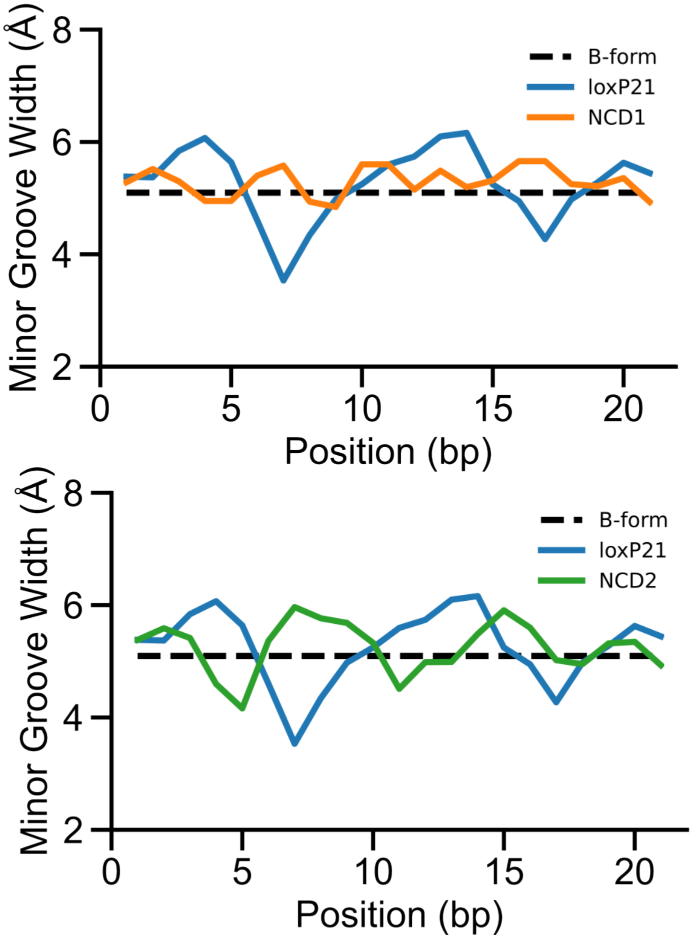


**Supplemental Figure 2.** Predicted DNA minor groove width of cognate (loxP21, blue) and noncognate (NCD1, orange and NCD2, green) indicate unique structure for the cognate DNA sequence.

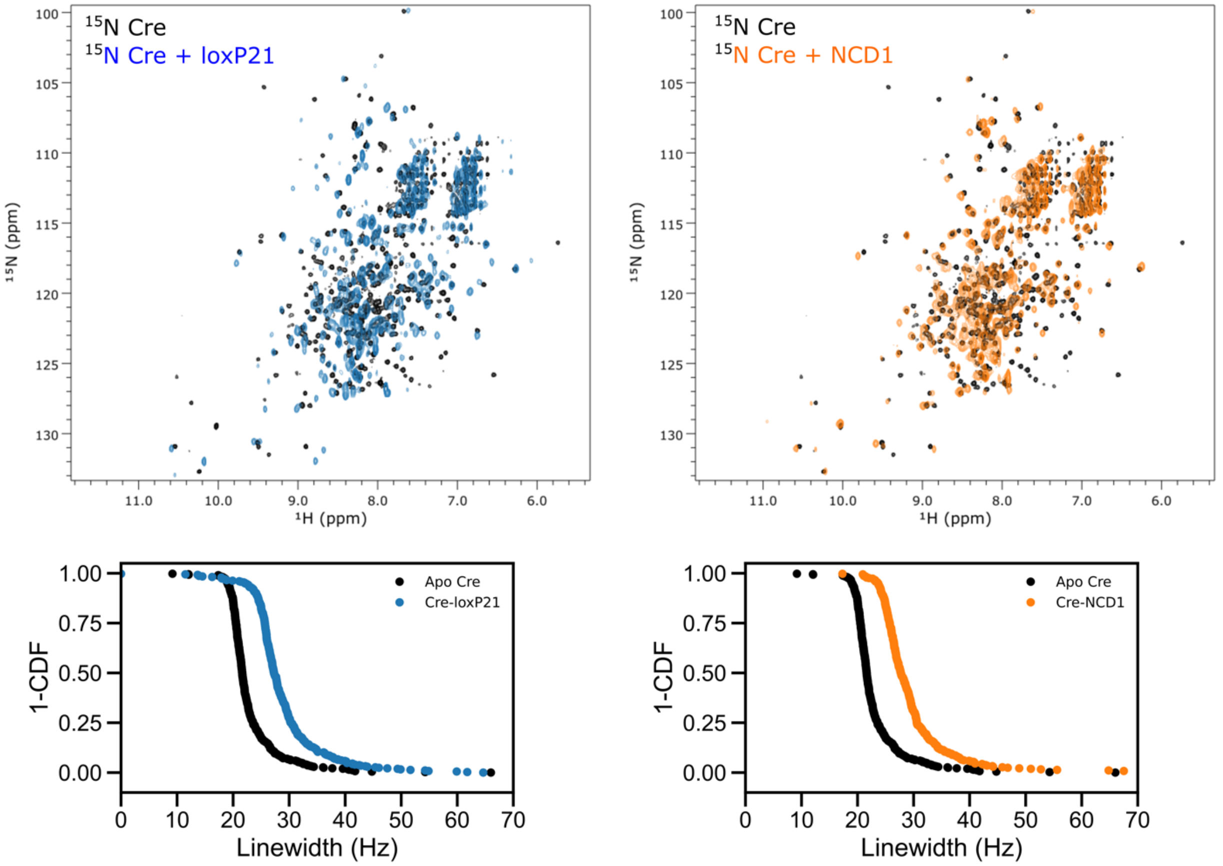


**Supplemental Figure 3.** TROSY HSQC Spectra of $^{15}$N Cre bound to either *loxP*21 or NCD1. For both *loxP*21 and NCD1, spectra of complexes show perturbations of chemical shifts and increases in $^{15}$N linewidths as a result of binding DNA substrates.

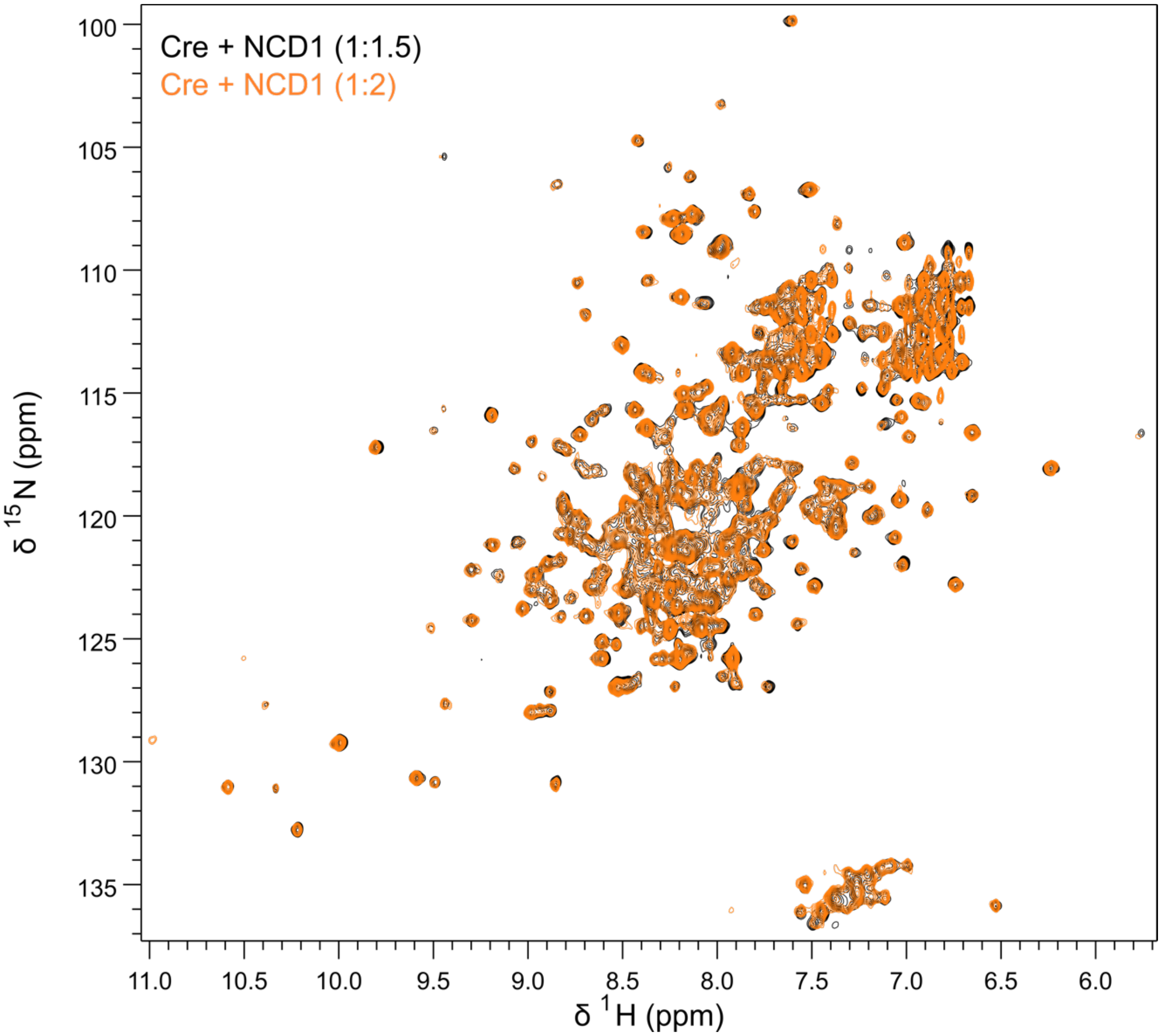


**Supplemental Figure 4.** TROSY-HSQC spectra of Cre bound to excess NCD1. TROSY-HSQC spectra were collected for Cre in complex with NCD1 at 1.5 or 2 molar equivalents of DNA. No significant chemical shift differences were observed in the protein indicating near complete binding to non-cognate DNA under these conditions.

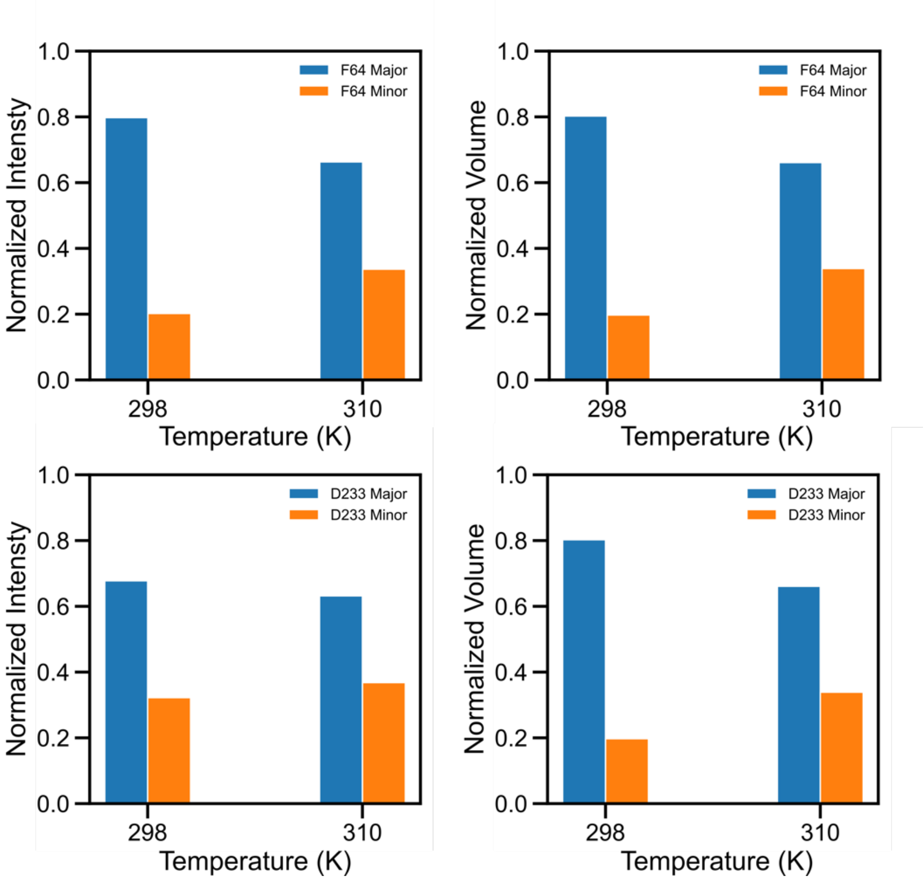


**Supplemental Figure 5.** Effect of temperature on populations for species with observed peak doubling. Increasing temperature from 298K to 310K results in an increase in peak intensity and volume for the minor species with a coincident decrease in the major species.

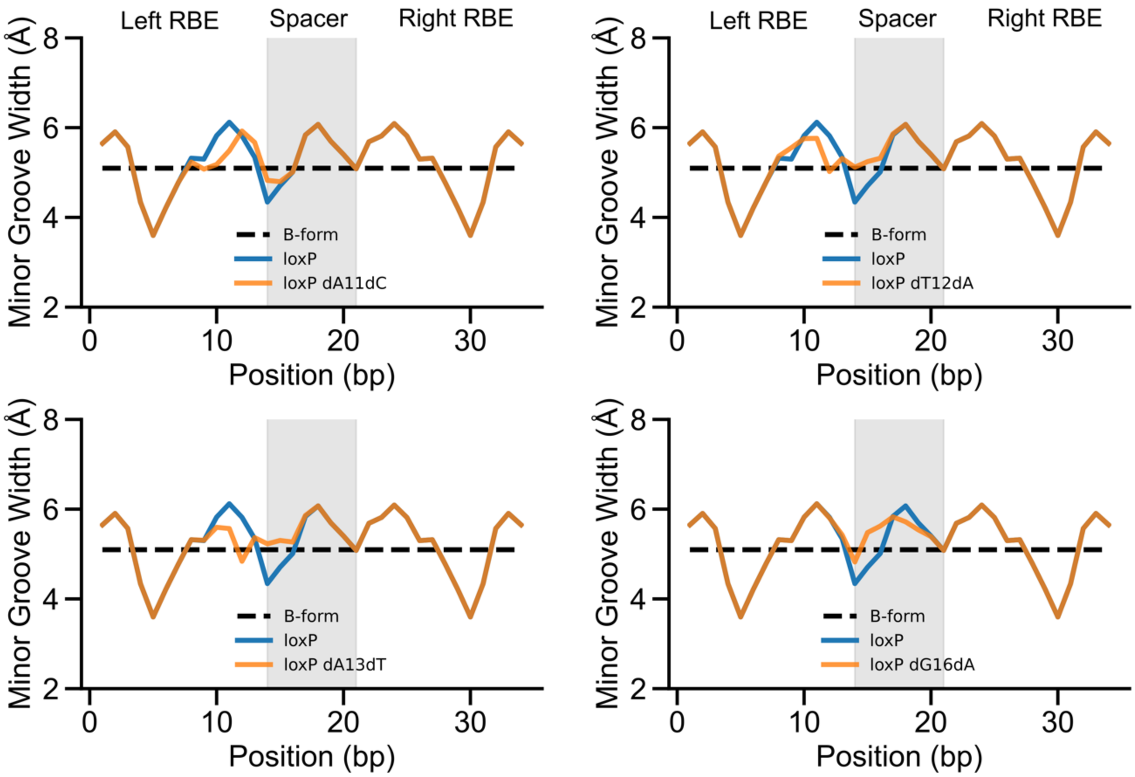


**Supplemental Figure 6.** DNA shape predictions comparing predicted minor groove widths for *loxP* and *lox* mutations that reduce recombination efficiency. Grey highlighted regions indicate the spacer.
